# MCH enables synchronized firing in the hippocamposeptal circuit to facilitate spatial memory

**DOI:** 10.1101/2020.05.25.114389

**Authors:** Jing-Jing Liu, Richard W. Tsien, Zhiping P. Pang

**Author notes:** Correspondence: Zhiping P. Pang.

## Abstract

Neuropeptide melanin-concentrating hormone (MCH) plays important roles in the brain including control of energy homeostasis, sleep, learning and memory. However, the synaptic and circuitry mechanisms underlying MCH-mediated regulations remain largely unknown. Here, we uncover that MCH modulates the hippocampo (HP) −dorsal lateral septum (dLS) −lateral hypothalamus neural circuit to facilitate spatial learning and memory. MCH achieves this function by enhancing both excitatory and inhibitory synaptic transmission via presynaptic mechanisms. The dLS neuronal spiking activity in response to HP CA3 excitatory inputs is strongly controlled by feed-forward inhibition (FFI) mediated by both GABA_A_ and GABA_B_ receptors. Endogenous MCH signaling enhances *Signal/Noise (S/N)* ratio of dLS neurons by increase the excitatory strengths, meanwhile decrease the overall dLS excitability by enhance inhibition which reduces dLS FFI, and consequentially enables dLS neurons to fire with high fidelity with HP synaptic inputs. Our data unravel the multifaceted synaptic mechanisms of MCH in the defined HP-dLS circuitry which may contribute to learning and memory.

## Introduction

Information flow mediated by synaptic transmission in the brain controls complex behavior and is exquisitely regulated by neuromodulators such as neuropeptides ^1, 2^. Melanin-concentrating hormone (MCH) is a hypothalamus-derived polypeptide that controls energy homeostasis ^3^, learning and memory ^4^, and social interactions ^5^. MCH-producing neurons (MCH neurons) are known to localize in the lateral hypothalamic area (LHA) as well as zona incerta that project to various brain regions including the hippocampus and the dorsal lateral septum (dLS) ^6-8^. MCH neurons are capable of releasing glutamate ^9^, γ-amino butyric acid (GABA) ^10^, and also producing other neuropeptides like nesfatin ^11^ and cocaine-amphetamine-regulated transcript (CART) ^12^. Activation, inhibition, or ablation of MCH neurons revealed that they play pivotal roles in reward ^13^, foraging behavior ^14^, as well as rapid eye movement (REM) sleep ^10, 15^. A recent study suggests that REM sleep-active MCH neurons are involved in forgetting hippocampus (HP)-dependent memories, likely by modulating HP neuronal activity ^16^. It is likely that MCH neurons exert their functions in both MCH-dependent and -independent manners ^16-23^, and therefore a critical mechanism underlying MCH regulation of hippocampal dependent memory storage remains unknown. system.

The dLS receives dense innervation relative to other brain regions from MCH neurons ^6, 7, 9^. The dLS functions as an important nodal point for information integration with strong inputs from various brain regions including the HP ^24, 25^ and the hypothalamus ^9, 24, 25^, and plays essential roles in social interactions ^26^, feeding ^27^, learning, and memory ^28, 29^. On the level of neuronal information coding, increases in spike rate- and phase-coupling are known to be essential in the HP-dLS circuit for place mapping, learning, and memory ^30-35^. A recent study showed that hippocampal-dependent cognition and neuronal encoding of spatial information may be transformed into firing patterns of dLS neurons, a potential route for the transformation of the ‘cognitive map’ into action ^28^. Neurons in the dLS also send projections to the LHA ^24, 25^, which is a brain region that governs motivational behavior ^36^. Yet understanding how endogenous MCH regulates the circuitry function of the HP-dLS-LHA remains unexplored.

Here, using a combination of genetics-aided neural tracing and systematic synaptic physiology, we uncovered a previously unknown and robust regulatory mechanism of endogenous MCH in the HP-dLS-LHA neural circuit. MCH increases signal to noise (*S/N*) ratio by suppressing the spontaneous firing of dLS neurons via enhancing GABA release into the dLS which suppresses the overall excitability of dLS neurons. Meanwhile, MCH facilitating HP CA3-to-dLS excitatory strength in the targeted neurons. The dLS neuronal spiking activity in response to hippocampal CA3 excitatory inputs is strongly controlled by feed-forward inhibition (FFI) mediated by both GABA_A_ and GABA_B_ receptors. The differential time courses of GABA_A_ and GABA_B_ receptor-mediated responses cast different inhibitory effects on postsynaptic neuronal firings. MCH in the dLS relieves the restraint on firing rate by dampening collateral FFIs via decreasing the overall excitability of dLS neurons. With facilitated upstream HP excitatory input, excitatory/inhibitory (E/I) balance thus is shifted to enable dLS neurons to fire in rhythm with high frequency dorsal hippocampal activity. We postulate that MCH regulates the HP-dLS circuitry via the aforementioned synaptic mechanisms to facilitate spatial memory

## Results

### Endogenous MCH regulates spontaneous activity of dLS-to-LHA projecting neurons

To specifically interrogate MCH neurons, AAV-DIO-Channelrhodopsin2 (ChR2)-eYFP was injected into the LHA of *pMCH*-Cre BAC transgenic mice ^10, 17^. Additionally, microfluorescent Retrobeads were injected into the LHA of the same animal to isolate dLS-to-LHA projecting neurons (**Fig. 1a**). Optogenetic stimulation reliably activated viral-infected neurons at up to 20Hz (**Fig. 1b&c**), which is within the physiological firing spectrum of MCH neurons *in vivo* during explorative behaviors (10∼40 Hz) in freely moving mice ^37, 38^. MCH neurons were found to innervate widely in the brain ^7^, with dense, varicose-like projections targeting the dLS (**Fig. 1d&e**). Consistent with a previous report ^9^, optogenetic activation of MCH axons induced excitatory postsynaptic currents (oEPSCs) mediated by glutamate release in dLS neurons (35 out of 115 neurons recorded) (**Fig. 1f**). These MCH neuronal oEPSCs in dLS-to-LHA projecting neurons consistently showed fast depletion during trains of stimuli (≥ 5 Hz) (**Fig. 1f&g**).

**Fig. 1.**
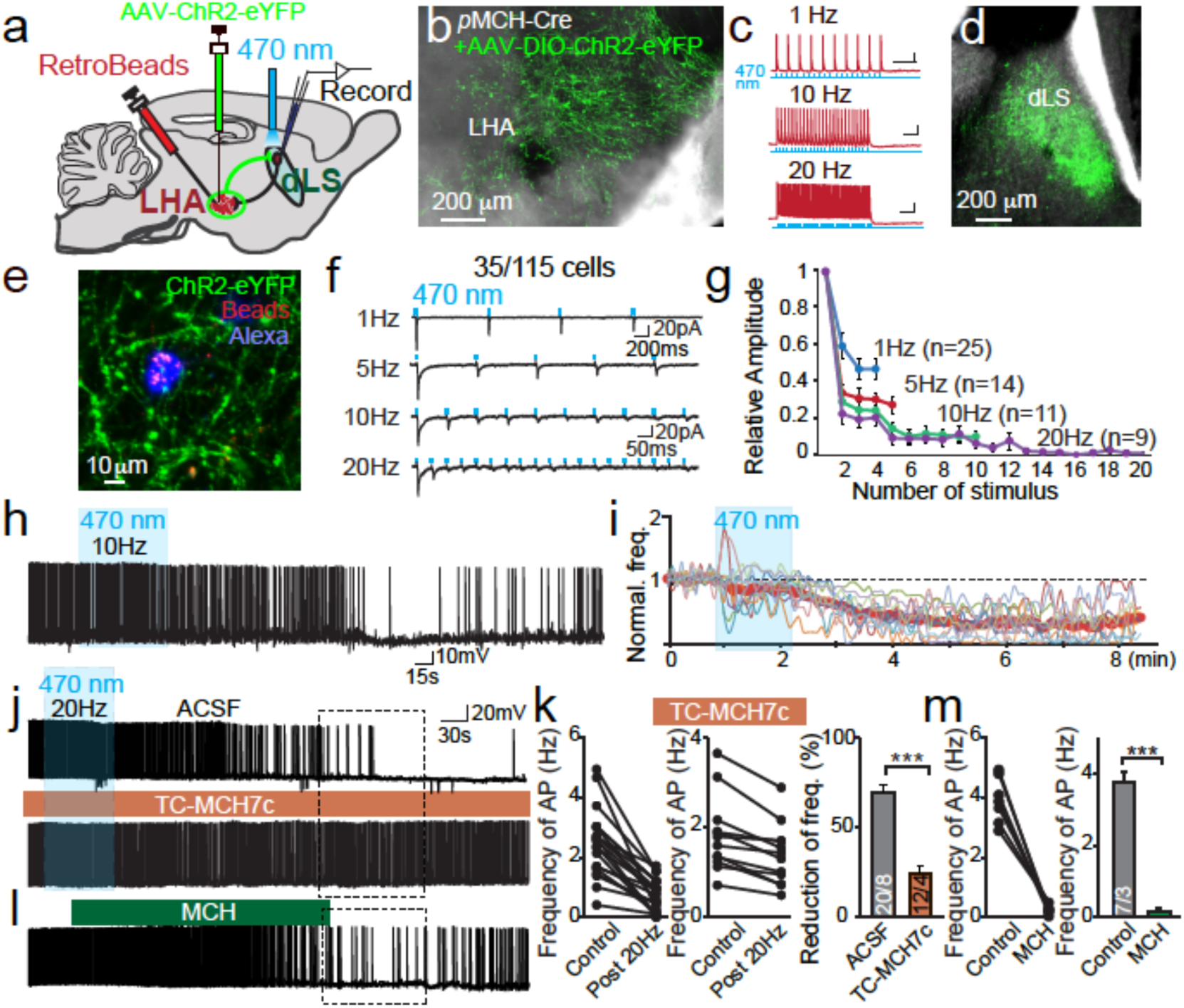
Endogenous MCH suppresses neuronal spontaneous activity of the dLS. **a**, Diagram representing the experimental setup. Retrograde fluorescent microbeads and AAV-DIO-ChR2 were sequentially injected into the LHA of *p*MCH-Cre mice. Retro-labeled dLS neurons were analyzed by whole cell patch clamp recording. **b**, Representative image of MCH-expressing neurons by expressing Cre-dependent ChR2-eYFP. **c**, LHA MCH-expressing neurons can be reliably evoked to fire action potentials up to 20Hz following optogenetic activation. **d**, ChR2-eYFP positive axonal projections from MCH neurons were identified in the dLS. **e**, An example of a Retrobead-labeled dLS-to-LHA projecting neuron that was recorded and loaded with Alexa Fluor 633 dye. **f**, Sample traces of MCH neuronal axonal optogenetic stimulation induced oEPSCs in dLS-to-LHA projecting neurons. 35 out of 115 cells showed oEPSCs. **g**, Pooled data showing that MCH neuronal origin oEPSCs depressed quickly. **h**, Sample trace showing that spontaneous action potential firing (sAP) of dLS neurons was suppressed by prolonged light stimulation (10Hz for 80s) on ChR-expressing MCH axons. **i**, Normalized frequency of sAPs in dLS-to-LHA projecting neurons. The thick red line shows the average frequency over time. **j**, Sample traces of sAPs under different conditions. *Upper panel*: 20Hz optical stimulation for 80s; *Lower panel:* recording performed in the presence of MCHR1 blocker, TC-MCH7c; **k**, Pooled data showing the changes of sAPs with or without MCHR-1 blocker. **l**, Exogenously applied MCH mimics the effect of optogenetic stimulation. **m**, Pooled data showing the frequency of sAPs after the application of MCH (600 nM). Data are mean ± SEM; numbers of neurons/animals analyzed are indicated in bars. Paired two-tailed Student’s t-tests were used: ***p<0.001.

The dLS is largely composed of GABAergic projecting neurons, which also form an extensive yet confined recurrent axonal plexus within the dLS region, an anatomical substrate for local collateral inhibition and feed-forward inhibition (FFI) ^39-42^. Indeed, the MCH neuron-derived oEPSCs were frequently followed by secondary inhibitory postsynaptic currents (oFFI-IPSCs) with a delay of a few milliseconds (**Supplementary Fig 1a&b**). Therefore, during a short-term stimulation (10 s; 10 Hz or 20 Hz), the overall acute impact of MCH nerve terminal activation on dLS neuronal firing varied, depending on the net gain of excitation (i.e. the oEPSCs) and inhibition (i.e. the oFFI-IPSCs) (**Supplementary Fig. 1c-f**).

To induce endogenous MCH neuropeptide release, we used prolonged high frequency stimulations ^43^, during which glutamate release from MCH fibers is likely minimal due to fast depletion (**Fig. 1f&g**). We first examined the spontaneous activity of dLS neurons and found that in neurons with spontaneous action potential firing (sAP) (28 out of 82 neurons recorded, 2.36±0.33 Hz), 10 Hz or 20 Hz 80-second optogenetic stimulations of MCH axon terminals led to a delayed suppression of the sAP activity in dLS neurons (**Fig. 1h-k, Supplementary Fig. 1g&h**). This suppressive effect is likely mediated by the MCH receptor-1 (MCHR1), as it was significantly blunted by application of Tc-MCH7c, a specific MCHR1 antagonist (**Fig. 1j&k**). MCHR1 is a G-protein coupled receptor (GPCR) and the only identified functional receptor expressed in the rodent brain ^44^ including the dLS (Fig. XX). Moreover, exogenous application of MCH mimicked the effect observed with optogenetic activation of MCH fibers in the dLS neurons (**Fig. 1l-m**). These data suggest that endogenous MCH release can regulate neuronal excitability in the dLS neurons that are projecting to the LHA. Nevertheless, it has been reported that the dLS neurons rarely express receptors for MCH ^45-47^, which precludes the direct effect of MCH signaling on these neurons but favors a possible regulation via synaptic transmission.

### Endogenous MCH facilitates hippocampal CA3-to-dLS excitatory synaptic inputs

Because a suppression of excitatory synaptic strengths could account for MCH suppression of sAPs, we next investigated the impact of endogenous MCH signaling on excitatory synaptic transmission in the dLS. Both spontaneous and evoked excitatory postsynaptic currents (sEPSCs and eEPSCs), isolated pharmacologically by application of picrotoxin (PTX), were examined in dLS-to-LHA neurons in the absence or presence of MCH. Contrary to initial expectations, prolonged trains of optogenetic stimulation of MCH fibers enhanced both sEPSCs and eEPSCs in dLS-to-LHA projecting neurons (**Fig. 2a-c, Supplementary Fig. 2a-c**). This augmentation of excitatory synaptic transmission is likely the result of an increase in presynaptic release probability of glutamate due to: ***1)*** an increase in the frequency of sEPSCs (**Supplementary Fig. 2a-c**); ***2)*** augmentation of the amplitudes in both AMPA receptor (AMPAR)- and NMDAR-eEPSCs with no change in the AMPAR-eEPSC/NMDAR-eEPSC ratio (**Fig. 2a&b**); and ***3)*** a decrease in the paired-pulse ratio (PPR) (**Fig. 2c&d**). Interestingly, this observed impact of optogenetic manipulations on EPSCs was fully abolished in the presence of two different MCHR1 blockers, TC-MCH7c or SNAP94847 (**Fig. 2c&e, Supplementary Fig. 2d-g**). This strongly indicate that MCHR1 signaling is active in the dLS, likely located on the presynaptic terminals of synaptic inputs into the dLS. To further support this, exogenous MCH or its analog [Ala17]-MCH showed a consistent augmenting effect on sEPSCs in these dLS-to-LHA projecting neurons (**Supplementary Fig. 2h-m**).

**Fig. 2.**
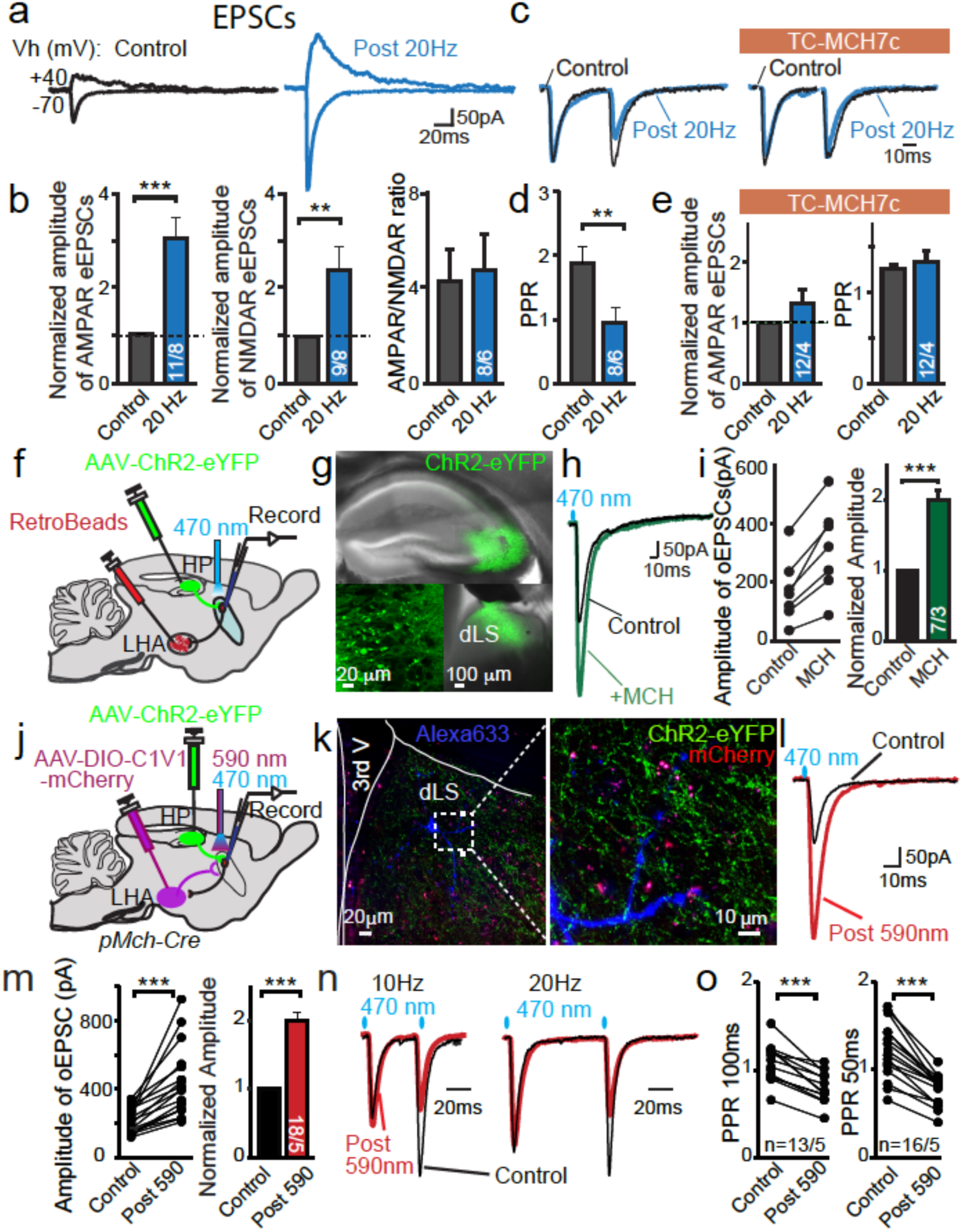
MCH enhances excitatory synaptic transmission in the hippocampus-to-dLS pathway. **a**, Sample traces of evoked AMPAR- (Vh=-70 mV) and NMDAR- (Vh=+40 mV) mediated EPSCs in dLS-LHA neurons by field stimulations before (control) and after 20Hz 2min optogenetic stimulation of MCH axons expressing ChR2. **b**, Pooled normalized data showing the amplitude of AMPAR-EPSCs, NMDAR-EPSCs and ratios of AMPAR-EPSC/NMDAR-EPSC. **c**, Superimposed sample traces of evoked EPSCs by paired pulses of field stimulations pre and post prolonged optogenetic stimulation in the absence or presence of MCHR-1 antagonist TC-MCH7c. **d**, Pooled data of paired pulse ratio (PPR) and the normalized amplitudes of evoked EPSCs. **e**, Normalized AMPAR-EPSCs and pooled data of PPRs for AMPAR-EPSCs in the presence of TC-MCH7c. **f**, Diagram representing the experimental setup to functionally illustrate the connections of the hippocampo-septo-LHA pathway. RetroBeads were injected into the LHA while AAV-ChR2-eYFP was injected into the ventral HP. Retro-labeled dLS-LHA neurons were analyzed by whole cell patch clamp recording. **g**, Viral mediated expression of ChR2-eYFP in the CA3 neurons. The inserts below show CA3 neurons expressing ChR2-eYFP and ChR2-eYFP labeled axonal projections in the dLS. **h**, Sample traces of optogenetically stimulated EPSCs (oEPSCs) recorded from dLS-LHA neurons, pre (black trace) and post (green trace) bath application of MCHR agonist MCH (600nM). **i**, Pooled data of oEPSCs. **j**, Diagram representing the experimental setup of dual optogenetic stimulations of hippocampal origin for oEPSCs by ChR2 (470nm) and of MCH neuronal origin for endogenous release of MCH peptide by red shifted C1V1 (590nm). AAV-ChR2-eYFP was injected into the ventral HP, while AAV-DIO-C1V1-mCherry was injected into the LHA of *p*MCH-Cre mice. dLS neurons were analyzed by whole cell recording. **k**, Viral mediated expression of C1V1-mCherry in MCH neurons and ChR2-eYFP in hippocampal CA3 neurons. Infected neuronal fibers from the HP (eYFP+) and MCH neurons (mCherry+) with a recorded neuron loaded with Alexa633 in the dLS region. **l**, Optogenetics-induced oEPSCs in the dLS pre and post prolonged 590nm light activation made on C1V1 positive MCH axons. **m**, Pooled data of amplitudes of single oEPSCs. **n**, Sample traces of oEPSCs after paired-pulse optogenetic stimulation. **o**, Pooled data of PPRs of oEPSCs at stimulation intervals of 50 and 100 ms. Data are mean ± SEM; numbers of neurons/animals analyzed are indicated in bars. Paired two-tailed Student’s t-tests were used: * p<0.05, ** p<0.01, ***p<0.001.

It has been known that the major excitatory inputs to the dLS are from the hippocampal formation ^24, 48^. In particular, the hippocampal CA2 projections are mainly restricted to the midline subsections of the dLS, which were recently found to be involved in social aggression ^26^ and fear conditioning ^49^; meanwhile the CA3 projections are confined to the lateral subsections of the dLS (**Fig. 2f&g, Supplementary Fig. 3a**). Coincidentally, hippocampal pyramidal neurons express high levels of MCHR1 mRNA ^47^. Thus, it is conceivable that MCH modulates HP-to-dLS projections. Indeed, MCH doubled the HP-to-dLS excitatory synaptic strengths in dLS-to-LHA projecting neurons (**Fig. 2h&i, Supplementary Fig. 3c-e**), accompanied by a decrease in PPRs (**Supplementary Fig. 3f**). Next, in order to directly address whether endogenous MCH has a similar effect, we took advantage of dual-color optogenetics ^50^. In each *pMCH*-Cre animal, we infected MCH neurons with Cre-dependent AAV-C1V1-mCherry, a red spectral (590 nm) shifted ChR, and simultaneously expressed ChR2-eYFP in the CA3 region (**Fig. 2j, Supplementary Fig. 3g-i**). Both eYFP-positive terminals (presumably originating from the HP) and mCherry-positive fibers (from MCH neurons in the LHA) were observed in the proximity of dendritic arborization of the dLS neurons, from which the recordings were performed (**Fig. 2k**). Remarkably, similar to that observed after exogenous application of MCH (**Fig. 2h&i**), the amplitude of HP-to-dLS oEPSCs evoked by 470 nm light nearly doubled after optogenetic activation of the MCH fibers using 590 nm light (2 min at 20 Hz) (**Fig. 2l&m**). Once again, the PPR significantly decreased (**Fig. 2n&o**), suggesting an involvement of presynaptic regulation. The augmentation of the HP-to-dLS oEPSCs after 590 nm activation of MCH fibers could be blocked through the use of two different MCH1R blockers, TC-MCH7c or SNAP94847 (**Supplementary Fig. 4**).

Collectively, although these data could not explain the underlying mechanism of MCH-mediated suppression of sAP (**Fig. 1h-m**), yet they strongly suggested that endogenous MCH strengthens information mediated by HP-to-dLS excitatory synaptic inputs via a presynaptic mechanism.

### Regulation of inhibitory synaptic transmission by MCH in dLS-to-LHA projecting neurons

Since the balance of *E/I* synaptic transmission could account for the final outcome of neuronal excitability and signal integration, we next explored how MCH signaling may regulate inhibitory synaptic transmission in the dLS neurons that project to the LHA. Inhibitory postsynaptic currents (IPSCs) were pharmacologically isolated by application of CNQX and D-APV, which block excitatory synaptic transmission. We found the frequency of spontaneous IPSCs (sIPSCs) drastically increased after prolonged optogenetic activation of MCH fibers in the dLS (**Fig. 3a&b, Supplementary Fig. 5a&b**). The amplitude of evoked IPSCs (eIPSCs) also increased, which was accompanied by a decrease in the PPR (**Fig. 3c&d**). Similar to that of EPSCs, the impact on inhibitory transmission from optogenetic activation of MCH nerve terminals was fully abolished by MCHR1 blockers (**Fig. 3c&d, Supplementary Fig. 5c&d**), but mimicked by the application of exogenous MCH (**Supplementary Fig. 5e&f**). Since dLS neurons rarely express MCHR1 ^45-47^ (Supplementary Figure xxxx), this suggests that MCH likely acts on the inhibitory input(s) from outside of the dLS ^41^.

**Fig. 3.**
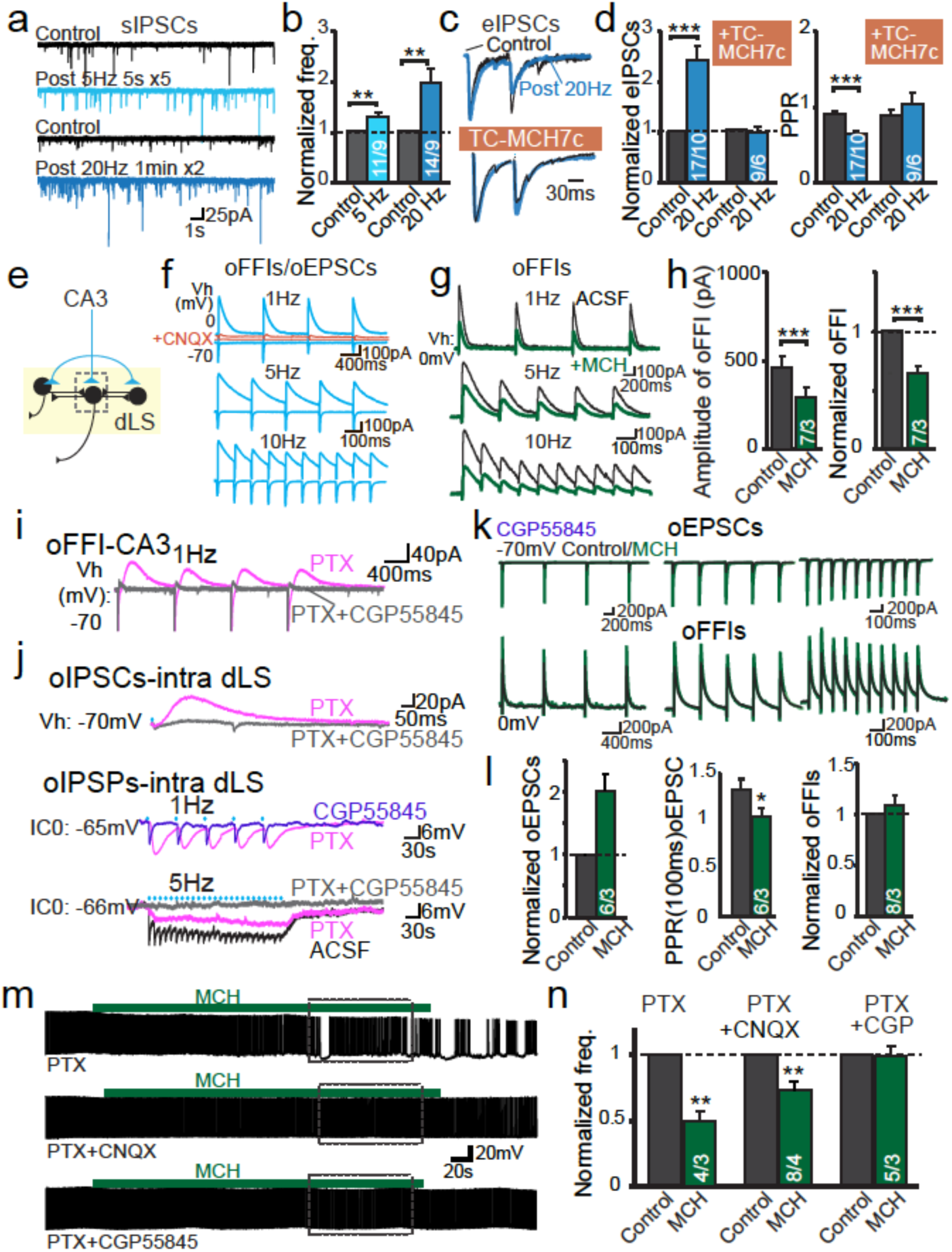
Augmentation of inhibitory synaptic transmissions by MCH accounts for the suppressive effect on dLS neuronal activity and excitability. **a**, Sample traces of spontaneous IPSCs (sIPSCs) mediated by GABA_A_ receptor (GABA_A_R) in dLS-to-LHA projecting neurons, pre and post 20Hz, 2min optogenetic stimulation of ChR-positive MCH axons. AMPAR blocker CNQX and NMDAR blocker AP5 were present in the bath for the isolation of inhibitory transmission. **b**, Pooled data showing the facilitation of sIPSCs. **c**, Superimposed sample traces of evoked IPSCs by paired pulses of field stimulations (eIPSCs) pre and post prolonged optogenetic stimulation, with or without the presence of MCHR antagonist TC-MCH7c in bath solution. **d**, Pooled data of normalized amplitudes of eIPSCs and PPRs. **e**, Diagram depicts hippocampal input-induced oEPSCs and feed forward inhibition (oFFI). AAV-ChR2-eYFP was injected into the ventral HP. **f**, Sample traces of primary HP-origin oEPSCs and secondary oFFIs in dLS neurons at varied frequencies. Both components were blocked in the presence of CNQX. **g**, Sample traces of hippocampal oFFIs pre (black traces) or post MCH (green traces). **h**, Pooled data. **i**, Sample traces of the hippocampal oEPSCs and GABA_B_R-mediated oFFI-IPSCs. **j**, Sample traces of the sum and isolation of both GABA_A_R- and GABA_B_R-mediated components in collateral oIPSCs and oIPSPs. **k**, Sample traces of primary HP-origin oEPSCs and secondary oFFIs in dLS neurons in the presence of GABA_B_R antagonist CGP55845 at varied frequencies pre (black traces) or post MCH (green traces). **l**, Pooled data showing the normalized oEPSCs, PPR and normalized oFFIs in the presence of CGP55845. **m**, Sample traces showing the impacts of MCH over sAP of dLS neurons during the blockage of GABA_A_R, GABA_A_R and AMPAR, or GABA_A_R and GABA_B_R. **n**, Pooled data of the amplitudes and PPRs of oEPSCs. *Right panel:* Pooled data of the amplitudes of oFFIs. Data are mean ± SEM; numbers of neurons/animals analyzed are indicated in bars. Paired two-tailed Student’s t-tests were used: * p<0.05, ** p<0.01, ***p<0.001.

The enhanced inhibitory inputs in the dLS induced by MCH should decrease the excitability of dLS neurons, thus reducing the collateral synaptic strength. To address this hypothesis, we expressed ChR2 sparsely in the dLS and recorded light-evoked intra-dLS IPSCs (oIPSCs) in neighboring non-infected dLS neurons (**Supplementary Fig. 6a**). Indeed, we found that the intra-dLS oIPSCs were suppressed following MCH application (**Supplementary Fig. 6b&c**). Somewhat to our surprise, MCH also suppressed the ChR2-evoked HP-to-dLS oEPSCs induced FFIs (i.e. oFFIs) (**Fig. 3e-h**), despite oEPSCs enhancement by MCH (**Fig. 2f-o**). These data suggest that the undermined collateral inhibition within the dLS mediated by MCH is likely due to the elevated GABA tone (from inputs outside the dLS) which outpowers the augmentation of excitatory synaptic inputs, both affected by MCH. Therefore, the excitation/inhibition (E/I) mediated by excitatory glutamatergic inputs and inhibitory GABA_A_ and GABA_B_ signaling determines either the neurons would be overall excited or inhibited. This hypothesis was strengthened by the observation of GABA_B_ receptor (GABA_B_R)-mediated slow collateral synaptic inhibition (time-to-peak of 207.8±5.9 ms) in the dLS, in addition to GABA_A_ receptor (GABA_A_R)-mediated fast inhibition (**Fig. 3i&j, Supplementary Fig. 7**). In support, high expression levels of GABA_B_R were reported in the dLS region ^51^. Interestingly, in the presence of CGP55845, MCH failed to undermine HP-dLS oFFIs, though it continuously strengthened excitatory input from the HP (**Fig. 3k&l**).

Collectively, these results indicate that MCH facilitates GABA release in the dLS, which results in an overall tonic suppression in the dLS neuronal excitability via GABA_A_R- and GABA_B_R-mediated signaling. We hypothesized that the overwhelmingly enhanced inhibition was the underpinning of sAP suppression by endogenous MCH in the dLS neurons, and this was confirmed by the result that in the presence of both PTX and CGP55845, MCH no longer suppressed sAP firing in the dLS neurons (**Fig. 3m&n**).

### MCH enables high-frequency-coded information flow

Given the multilevel synaptic regulatory effects by MCH in the dLD, we next sought to address how MCH modulates neuronal signal integration in the dLS. To this end, we specifically examined the generation of APs in dLS-to-LHA projecting neurons induced by HP inputs (**Supplementary Fig. 8**). During 1 Hz stimulation we observed relative high fidelity in stimulus-locked optogenetic excitatory postsynaptic potential (oEPSP)-induced AP generation in dLS neurons, with a success rate of 78.5± 5.13% (n=38) (**Supplementary Fig. 8**). Strikingly, as the stimulation frequency increased to 5, 10, and 20Hz, a reduction in oEPSP-AP production was observed. During 5Hz stimulation (25 light pulses in 5s), approximately 13 time-locked spikes were generated in dLS neurons, which correlates to a near 50% coding efficiency for the HP-dLS-LHA circuit. The efficiency further reduced to 45% at 10Hz and decreased as low as 20% at 20 Hz (**Supplementary Fig. 8**). We hypothesize that the slow and prolonged (∼200 ms) GABA_B_R-mediated hyperpolarization works as an empirical ‘*low pass filter’* to limit the transduction of higher frequency-encoded information from the HP to the dLS. Application of GABA_B_R blocker CGP55845 largely blocked the progressive failure of oEPSP-APs (**Supplementary Fig. 9a**).

The dLS cells that had 100% oEPSP-AP production following 1Hz stimulation (n=22) were selected for further pharmacological perturbation of MCH signaling (**Supplementary Fig. 9b**). As predicted, the application of MCH drastically elevated the success rate of oEPSP-AP production, over 95% at 10Hz and 80% at 20Hz (**Fig. 4a&b**). In the presence of MCH, both GABA_A_R- and GABA_B_R-mediated components in HP-to-dLS oFFI-inhibitory postsynaptic potentials (IPSPs) were significantly attenuated due to the suppression of collateral inhibition (**Supplementary Fig. 9c, Fig. 3e-h**). Through such fine synaptic regulation, MCH signaling allows the removal of the ‘*low pass filter’* function set by FFIs in the dLS neuronal network. Not surprisingly, GABA_B_R blockers occluded the effects of MCH (**Supplementary Fig. 9**).

**Fig. 4.**
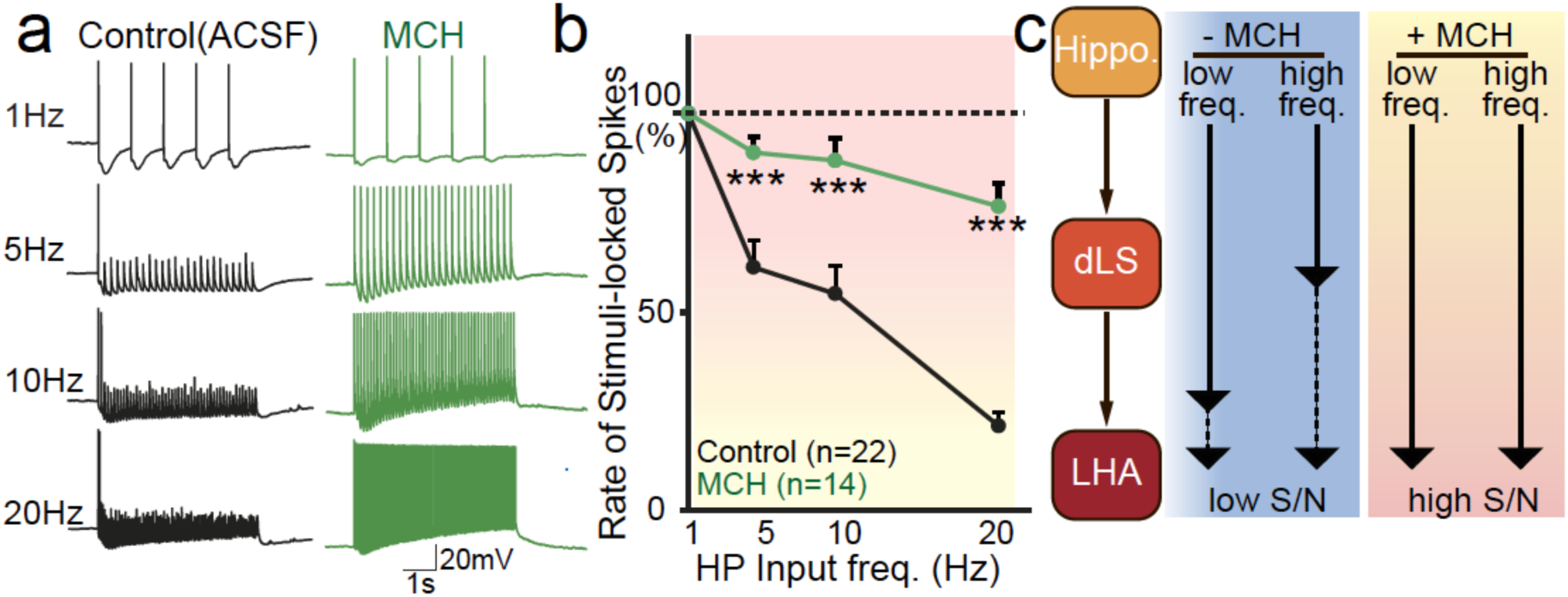
MCH enables high-frequency-coded information flow in the HP-dLS-LHA circuit. **a**, Sample traces of oEPSP-APs of dLS-to-LHA projection neurons under different conditions as labeled. **b**, Plot depicts the pooled data of the ratio of oEPSP-APs of dLS neurons as functions of the frequency of hippocampal optogenetic stimulations. **c**, Model of permissive effects of MCH in spike coupling between hippocampal inputs and dLS neuronal firing. All data are presented as mean ± SEM. Student’s t-tests were used: ***p<0.001.

Although the blockage of GABA_B_R signaling could increase the fidelity of postsynaptic oEPSP-APs, similar to that observed after MCH application, there are important differences between these two different manipulations. *First*, MCH affects both GABA_A_R and GABA_B_R-mediated FFI-IPSPs, but CGP55845 only affects the latter. *Second*, MCH increases the *S/N* ratio (excitatory inputs/spontaneous firings), whereas blockage of GABA_B_R signaling likely increases the overall baseline activity and excitability of dLS neurons, thus increasing both *S* (i.e. signal) and *N* (i.e. noise). *Third*, there is no known endogenous GABA_B_R antagonist in the brain. Collectively, our data strongly suggest that the dLS functions to compute and control information flow from the hippocampal CA3 to downstream targets, and concurrent MCH release enables high frequency encoded information to be integrated in the dLS (**Fig. 4c**) via multifaceted yet precise control of synaptic transmission.

The neuronal coding mechanism in the HP-dLS circuit has been intensively explored, with early findings demonstrating that the increase in spike rate and phase coupling are essential in this circuit for place mapping, learning, and memory ^30-35^. Most of the HP pyramidal cells fire at less than 2 Hz while at rest but increase their firing rate up to 50Hz (average 9∼12 Hz) when animals are in their place fields ^30, 31, 52^. Similarly, dLS neurons have also been suggested to contain place-related activity and a place map ^28^. Their AP firing rate increases up to above 20 Hz when animals are in their place field, and simultaneously show phase procession relative to HP theta waves ^28-30, 33, 53^. These evidence suggest that hippocampal-dependent cognition and neuronal encoding of spatial information may be transformed into firing patterns of dLS neurons, a potential route for the transformation of the ‘cognitive map’ into action ^28^.

### MCH-mediated signaling in the dLS affects spatial learning and memory

Given that MCH enables the high frequency coupling of HP input to dLS spikes, we hypothesized that dLS MCH signaling is involved in behaviors that are dependent on the HP-dLS circuitry ^54-56^. To test this hypothesis, MCH signaling was manipulated by pharmacological interventions, in which MCH agonist, antagonist, or vehicle controls were administered into the dLS region via bilateral cannulae^37^ (**Supplementary Fig. 10a&b**). The Morris Water Maze (MWM) test indicated that the MCH agonist increased spatial learning, whereas the antagonist impaired it. (**Fig. 5, Supplementary Fig. 10d**). Meanwhile, these manipulations showed no significant impact on RotaRod motor learning or locomotion (**Supplementary Fig. 10c&e**). In the MWM test, the MCH antagonist group performed significantly worse in almost all the parameters measured compared to the control or the MCH group, whereas the MCH group showed increased ability and precision in locating target areas relative to the control group. For example, mice in the MCH treatment group spent longer time searching through the quarter-quadrant that used to contain the platform, and their head entry time in the platform area was about twice as long as that of controls (**Fig. 5d-f**). Additionally, nesting behavior and social recognition also showed sensitivities to the perturbation of MCH signaling in the dLS, which were indicated to be under hippocampal septal control ^26, 57, 58^ (**Supplementary Fig. 10f&g**). Overall, these findings suggest that MCH signaling in the dLS is indispensable for hippocampal-dependent behaviors, in particular for accurate encoding and consolidation of precise space locations in working memory ^52, 59^.

**Fig. 5.**
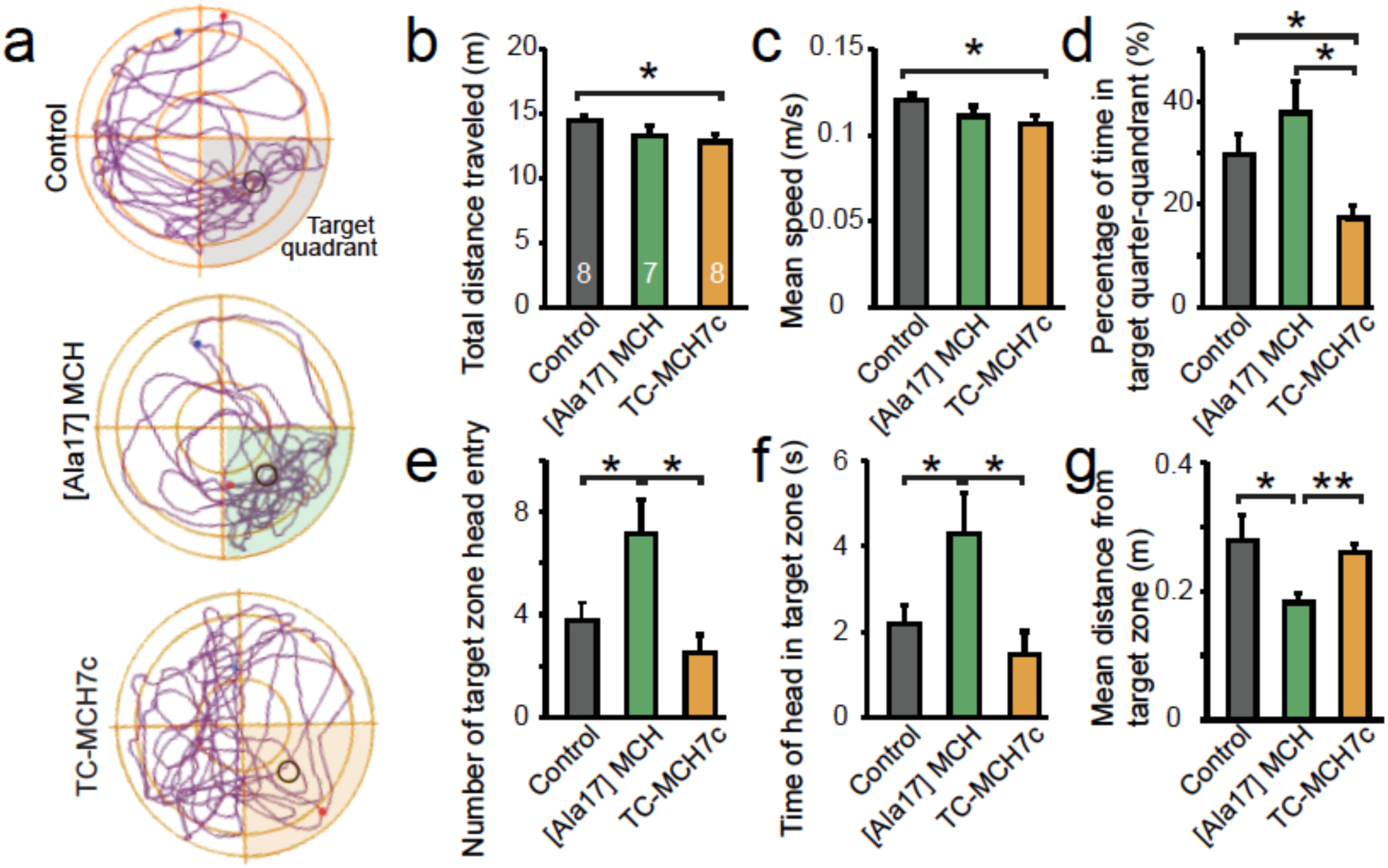
MCH is required for HP-dependent spatial learning and memory consolidation. **a**, Representative travel paths of animals in Morris Water Maze test. Animals were locally infused with vehicle (phosphate-buffered saline), MCHR agonist [Ala17]-MCH, or MCHR antagonist TC-MCH7c by bilateral cannula targeting the dLS. **b**, Pooled data of the total distance traveled. **c**, Mean speed. **d**, Percentage of time in target quarter-quadrant. **e**, Number of target zone head entries. **f**, Time of head in target zone. **g**, Mean distance from target zone. All data are presented as mean ± SEM; numbers animals analyzed are indicated in bars. ANOVA tests were used: * p<0.05, ** p<0.01, ***p<0.001.

## Discussion

In this study we systematically analyzed the complex synaptic and circuitry regulatory functions of MCH signaling in the CA3-dLS-LHA neurocircuit. We showed compelling evidence that: ***1)*** endogenous MCH augments both excitatory (including CA3 inputs) and inhibitory synaptic releases in the dLS via presynaptic mechanisms; ***2)*** GABA_B_ receptor-mediated slow inhibitory postsynaptic response has a profound impact on neuronal coding in the HP-to-dLS circuit; ***3)*** intra-dLS collateral FFIs also have a crucial impact on HP origin neuronal coding; through these mechanisms ***4)*** MCH increases the synchrony between HP CA3 firing and postsynaptic dLS neuronal firing at higher frequency which may contribute to spatial learning.

In addition to many important roles played by MCH peptide ^3-5^, MCH-producing neurons likely also function through molecules other than MCH. For example, ablation of MCH neurons and MCH knockout has different impacts on glucose metabolism^17-20^. Microinjection of MCH in the HP in rats facilitated memory retention in an inhibitory avoidance test ^21^. In support, knockout of MCHR1 led to cognitive deficits ^22^ and impaired long-term synaptic plasticity in the HP ^23^. However, a recent paper showed that activation of MCH neurons impaired memory, whereas ablation or inhibition of MCH neurons during REM sleep facilitated memory ^16^. Given that MCH neurons are capable of releasing glutamate ^9^, GABA ^10^, and producing other neuropeptides like nesfatin ^11^ and CART ^12^, these neurotransmitters and/or neuromodulators may serve varied physiological functions during MCH neuronal activity change. Indeed, the involvement of MCH release in the HP was not specifically addressed in this recent study during manipulations of MCH neuronal activity ^16^. Our data, on the other hand, suggest that dLS MCH signaling facilitates HP dependent spatial learning, likely via coupling of high frequency HP-dLS neuronal coding. We believe that a number of factors may contribute to the discrepancies in studies on MCH neuronal (or MCH) functions: ***1)*** MCH neurons contains neuroactive substances other than MCH ^9-12^ that may affect metabolism and memory differentially; ***2)*** MCH neurons potentially are molecularly diverse depending on their projection sites ^60^; ***3)*** MCH neurons project to different brain regions and may release a different set of neurotransmitters or neuromodulators in a specific downstream target, for example, release glutamate in the dLS ^9^, but release GABA in the tuberomammillary nucleus^10^; and ***4)*** MCH neuronal-mediated HP-dependent memory formation is upstream of dLS gating, and fine regulation of dLS coding by MCH actually enhances memory formation (Fig. 5). Clearly, further detailed molecular and circuitry mechanisms of MCH in various brain regions and different cell-subtypes are in need of elucidation.

FFI in neuronal networks controls input-output signal computation and neuronal coding ^61^. Our result indicates that collateral inhibition (substrate of FFI), especially the inhibition mediated by GABA_B_ receptors, is crucial for HP-to-dLS neuronal coding. In this system, high frequency firing of presynaptic HP CA3 neurons, instead of exciting postsynaptic dLS neurons, caused “shutdown” of postsynaptic neuronal firing because of the FFIs (**Fig. 4 &. Supplementary Fig. 8**). This is clearly dependent of the slower time course of the GABA_B_R-mediated synaptic inhibition. Neuropeptides such as MCH may enable flexible circuit information processing in the HP-dLS-LHA circuit via regulating FFIs. Nevertheless, the detailed computational model of this system still needs to be developed.

Taken together, our results provide important insights into the neuronal basis of endogenous MCH function in the HP-dLS-LHA neurocircuitry (**Fig. 4c**). Concurrent MCH release in the dLS modulates synaptic transmissions from the HP to enhance both rate- and phase-relations between the two brain regions (i.e. the HP and the dLS), as well as facilitating working memory associated with enhanced spatial accuracy guided by environmental cues ^24, 52, 59^. The study provides a general framework on the mechanism by which neuropeptides may work in concert with fast synaptic transmission to control the coupling of different brain regions and information flow^1, 2^. The modulatory effects of neuropeptide on various synaptic inputs (e.g. excitatory and inhibitory), different postsynaptic receptors with different kinetics (e.g. GABA_A_ and GABA_B_ receptors) should be considered in elucidating the physiological function of endogenous neuropeptides.

## Acknowledgements

We would like to thank Drs. Alexander Kusnecov, Baijuan Xia and Audrey Chang for their generous help in the behavioral analyses. We would also like to thank Drs. Arnold Rabson, Nicola Francis and Ms. Denise Robles of Rutgers, Dr. Long-Jun Wu of Mayo Clinic, and Dr. Pascal Kaeser of Harvard Medical School for their constructive suggestions. The Pang laboratory is supported by NIH R01 AA023797 and in part by the Robert Wood Johnson Foundation (grant #74260) to the Child Health Institute of New Jersey.

## Contributions

J.J. Liu and Z.P. Pang conceived the study. J.J. Liu performed the experiments and analyzed the data. R.W. Tsien provided constructive intellectual input to the overall study. All authors contributed to the final manuscript.

## Materials and methods

### Animals

All procedures involving mice were approved by the Rutgers Robert Wood Johnson Medical School Institutional Animal Care and Use Committee (IACUC). All the animals used in this study were 5-11 week-old males unless otherwise stated. The animals are bred in the facility at ∼22°C and 35%–55% humidity. The *pMCH-Cre* transgenic mice and wild-type mice (C57BL/6) were purchased from Jackson Laboratory. Mice were group housed except when otherwise stated and maintained on a 12-hour light/dark cycle (lights on at 6:00 A.M.) with food and water available ad libitum.

### Retrograde Labeling

Mice (8 weeks old) were anesthetized using isoflurane and placed in a stereotactic frame (KOPF M1900). Standard serological and injection procedures were followed as described previously^62^. Red or green RetroBeads (100 nl, LumaFluor) were injected unilaterally into the dLS. Coordinates: Anteroposterior (AP): +0.5 mm, mediolateral (ML): ±0.45 mm, and dorsoventral (DV): -2.7 mm) or LHA (AP: -1.2 mm, ML: ±0.45 mm, and DV: -5.1 mm). Animals were allowed to recover in their home cages for 14 days to allow adequate retrograde transportation of the beads to the soma of input neurons. Injection sites were verified in all animals.

### Virus infection

The AAV-viruses used in this study include: AAV-EF1a-DIO-ChR2-YFP, rAAV2/CamKII-ChR2(H134R)-EYFP-WPRE-PA, and AVV-EF1a-DIO-C1V1(E122T/E162T)-TS-mCherry, and were purchased from the University of North Carolina Vector Core. The ChR2-YFP encodes a membrane-bound fusion protein, allowing visualization of both the cell bodies and axons of infected neurons. Viral-mediated protein expression was allowed for a period of 6 weeks prior to experimental manipulation. Injection sites were confirmed in all animals reported in this study. 0.6-1 μL virus was used for the injection of the Cre-dependent expression AAV virus. For the non-Cre dependent AAV virus, 0.2-0.4μL of virus was injected. The injection speed was 0.1 μL/10 min and the injection coordinates for HP were: AP: -1.6 mm from bregma; ML: ± 3.1mm; DV: -2.2 mm; for the LHA and dLS, same as the above mentioned. Six weeks post-surgery, mice were deeply anesthetized with Euthasol. Coronal dLS or LHA slices (300 μm) were prepared and whole cell patch clamp recordings were performed.

### In vitro Electrophysiology

Mice were anesthetized, decapitated, and brains were removed and quickly immersed in cold (4°C) oxygenated cutting solution (containing in mM: 50 sucrose, 2.5 KCl, 0.625 CaCl_2_, 1.2 MgCl_2_, 1.25 NaH_2_PO_4_, 25 NaHCO_3_, and 2.5 glucose, pH 7.3 with NaOH). Coronal hypothalamic or dLS slices, 300 µm in thickness, were cut using a vibratome (VT 1200S; Leica). Brain slices were collected in artificial cerebrospinal fluid (ACSF) and bubbled with 5% CO_2_ and 95% O_2_. The ACSF contained (in mM): 125 NaCl, 2.5 KCl, 2.5 CaCl_2_, 1.2 MgCl_2_, 1.25 NaH_2_PO_4_, 56 NaHCO_3_, and 2.5 glucose (pH 7.3 with NaOH). After at least 1 hour of recovery, slices were transferred to a recording chamber and constantly perfused with bath solution (30°C) at a flow rate of 2 ml/min. Whole cell patch clamp recordings were performed as described elsewhere ^13^. Briefly, patch pipettes with a resistance of 4∼6 MΩ were made from borosilicate glass (World Precision Instruments) with a PC-10 Narishige puller and filled with a pipette solution containing the following (in mM): 126 K-gluconate, 4 KCl, 10 HEPES, 4 Mg-ATP, 0.3 Na_2_-GTP, 10 phosphocreatine, (pH to 7.2 with KOH) or 90 K-gluconate, 40 CsCl, 1.8 NaCl, 1.7 MgCl_2_, 3.5 KCl, 0.05 EGTA, 10 HEPES, 2 Mg-ATP, 0.4 Na_2_-GTP, 10 phosphocreatine, 5 mM QX314 (pH to 7.2 with CsOH). Dextran Alexa Fluor 633 was included in the intracellular recording solution for cell labeling. After whole cell patch clamp was achieved, evoked EPSCs were recorded in voltage clamp at -70 or +40 mV in the presence of PTX (50 µM) to obtain specific AMPAR and NMDAR components, and IPSCs were recorded under voltage clamp at either -70 mV with CNQX (20 μM) or 0 mV according to the pipette solution used. For all field stimulation induced responses, the electrode was placed at least 125 µm laterally from the recorded neuron and a model 2100 Isolated Pulse Stimulator (A-M Systems) was used to generate stimulus. For the optical activation of ChR2-positive fibers, a LED light source was used to generate 470nm blue light with 1000 mA power and 1ms pulse width except when otherwise stated. Input resistance and series resistance were monitored throughout the experiments, and recordings were rejected if series resistance increased above 25 MΩ. Pharmacological agonists/antagonists were applied via perfusion into the bath recording solution after control recording. All data were sampled at 5 kHz and analyzed offline using ClampFit 10.2 (Molecular Devices, USA). For graphical representation, the stimulus artifacts were removed. MCH, TC-MCH 7c, SNAP 94847, [Ala)-MCH, Cart, PTX, TTX, D-APV, and QX-314 were obtained from Tocris Inc (USA); MCH (H-070-47) antibody was obtained from Phoenix Pharmaceutical.

### Experimental design and statistical analysis

The effect of MCH on synaptic transmission was evaluated before and after the bath application of 600 nM MCH in the same neurons that were recorded in dLS brain slices. Therefore, these experiments were not performed blindly. In all cases, three or more animals were used for each parameter. Individual sample sizes for slice patch clamp recording (n=number of neurons, labeled in each figures) are reported separately for each experiment. All data are presented as mean ± SEMs. Statistical comparisons before and after the application of MCH were made using paired two-tailed Student’s t-test. Statistical comparisons for different groups were made using unpaired two-tailed Student’s t-test.

### Intra-dLS cannula implantation and behavior tests

Cannulation was performed as previously described (1). Briefly, at six weeks of age, 30 male C57BL/6 wild-type mice were anesthetized and implanted with bilateral guide cannulas with dummy injectors (PlasticsOne) targeting the dLS region using the designated coordinates. Mice were individually housed for 5 weeks to allow recovery from the surgery, during which daily food intake and body weights were monitored.

All behavioral testing was conducted blindly using male mice. Behavior tests were arranged in order of least disruptive (absence of noxious stimulation: social interaction, open field/object recognition task, and RotaRod) to most disruptive (MWM, which involved physical stimuli, such as cool water). In order to reduce testing effects and fatigue, animals were given three weeks between the RotaRod and MWM. Each behavioral test was conducted between 6 AM to 12PM. 150 nL solution was infused into each side of the dLS per animal through an injector (with 0.2 mm extra in length) 30 min before daily training sessions to allow adaptation of the animals. After the MWM experiment, animals were perfused and implantation sites were confirmed.

RotaRod test consisted of three trials (with a 30min intertrial interval) per day over the course of 4 days. Each trial ended when a mouse fell off, or reached 300s. For the first 10 trials, the RotaRod accelerated from 4-40 revolutions/min in 300s. For the last 2 trials, the RotaRod accelerated from 4-40 revolutions/min in 150 s. The time to either fall off the rod or turn one full revolution was measured.

For nesting behavior, a new 5 cm x 5 cm square of pressed cotton material was placed in each cage every day after the completion of the daily RotaRod trial. The resulting nest was scored the next morning using a standard 5-point scoring method. The social interaction test was designed to analyze the amount of time a subject mouse spends in the side of a chamber with a familiar mouse confined in a wire cage, versus the other side of the chamber with a confined stranger mouse. The whole chamber was monitored and analyzed by Afasci Smartcage® system. The confining cages were made of stainless wire mesh, each fixed on the opposing walls. Five minutes were given to each experimental subject to explore and habituate to the environment with both wire cages left empty. Next, one random male mouse was introduced to the experimental subject while the other wire cage was left empty for 10min. Finally, the experimental subject was tested again with the mouse the subject met previously and another novel mouse (similar age and matching sex). Therefore, the dependent variable was the percentage of time spent within a given compartment. The open field/object recognition test addresses both non-specific and stimulus-specific exploratory behavior^63^. Subjects were placed in a 56×62 cm empty open field and allowed to explore the arena. Two sidewalls were patterned differently with strips or dots, whereas the other two sidewalls had no pattern. The locomotion and time spent in areas within the test field were recorded by a camera and analyzed by Anymaze software.

Morris water maze (MWM) task is a standard test of spatial learning and reference memory^64, 65^. As described here, an expedited form of the test was run over two days, with the first day consisting of 12 acquisition trials, while the second day involved a probe test. The MWM pool was 110cm in diameter and filled with water kept between 21° and 23°C. During the acquisition phase, a small, round platform was hidden 1–2 cm below the surface. The water was made opaque with non-toxic colored paint to ensure the platform was not visible. Permanent cues were located on a dark plastic curtain around the enclosed pool. The curtain served to hide the experimenter who operated the computer running the tracking software. In practice, the pool was divided into four quadrants, and each subject was randomly assigned a specific quadrant in which the platform was hidden for all trials. Each subject underwent twelve trials, a maximum of 60s each. The subject always started from a random quadrant (and always oriented to face the pool edge) that did not house the platform; therefore, for three consecutive trials the animal began in a different empty quadrant. If the subject found the platform within 60s, it was left to sit on the platform for five seconds and was then removed. If the subject did not find the platform at the end of the 60s trial, it was placed on the platform and allowed to sit for 15s before being removed. Latency to reach the platform was recorded for each subject in each trial. There was a 5-minute inter-trial interval between each of the twelve trials. Twenty-four hours later, the platform was removed and each subject underwent a full 120s probe trial. The percent time spent in the target quadrant where the platform was hidden during acquisition trials was recorded and used as the dependent variable for this probe test. A combination of tests, including t-tests, ANOVAs, chi-square tables of frequency, and correlations were used as appropriate.

## Supplementary Information

**Supplemental Figure 1.**
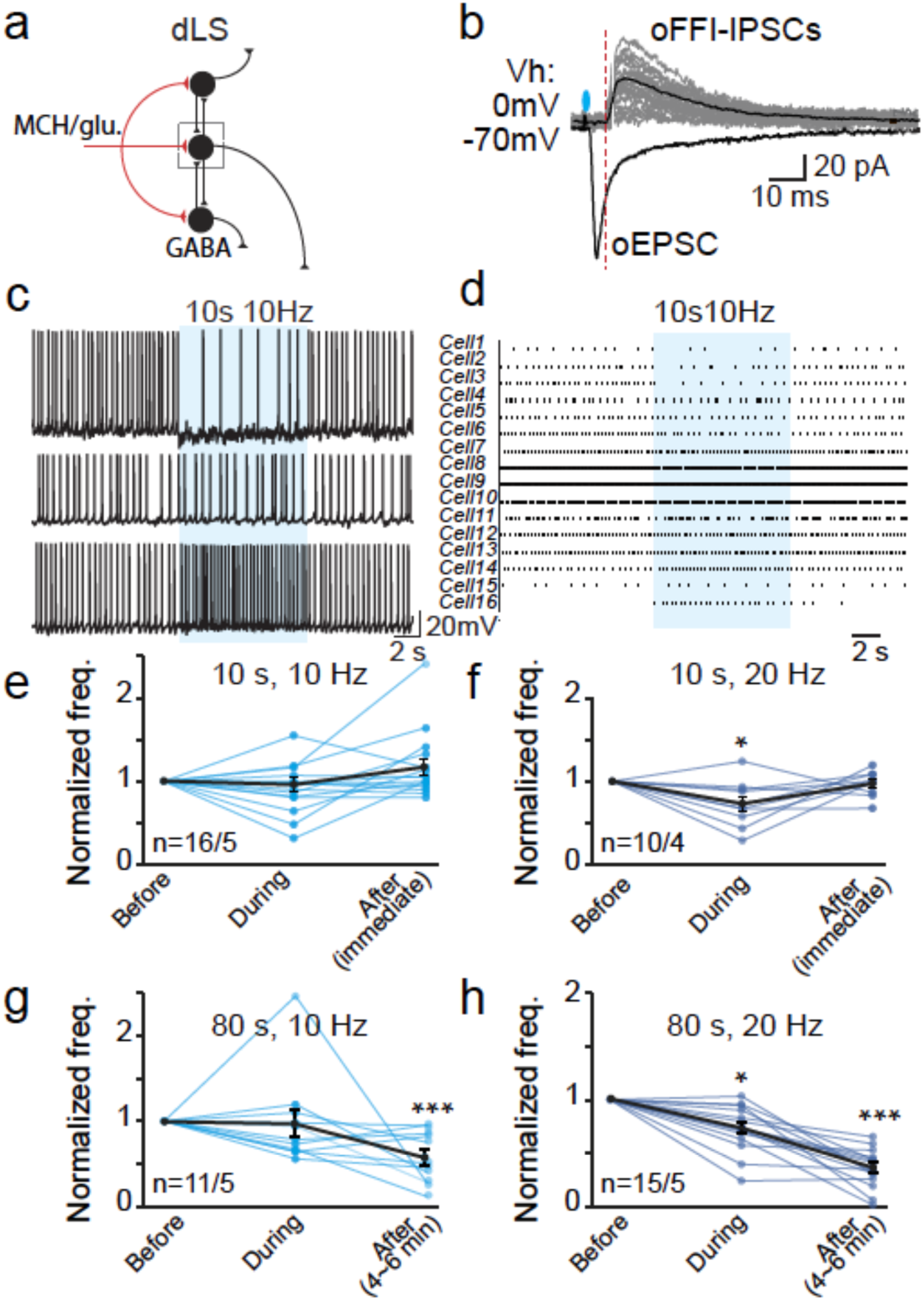
LHA MCH-dLS optogenetic stimulation may induce immediate influence on spontaneous dLS neuronal firing. **a**, Possible wiring diagram of dLS neurocircuitry. **b**, Sample traces of optogenetically-induced oEPSCs and feed-forward-inhibitory oIPSCs. **c**, Representative traces of dLS neuronal spontaneous action potentials (sAPs) before, during and after 10 s optogenetic stimulation of MCH fibers. **d**. Raster plot of sAPs. **e-h**, Normalized and pooled data of dLS neuronal firing under 10s or 80s, 10Hz or 20Hz stimulation. Data are mean ± SEM; numbers of neurons/animals analyzed are indicated. Paired two-tailed Student’s t-tests were used: * p<0.05, ** p<0.01, ***p<0.001.

**Supplemental Figure 2.**
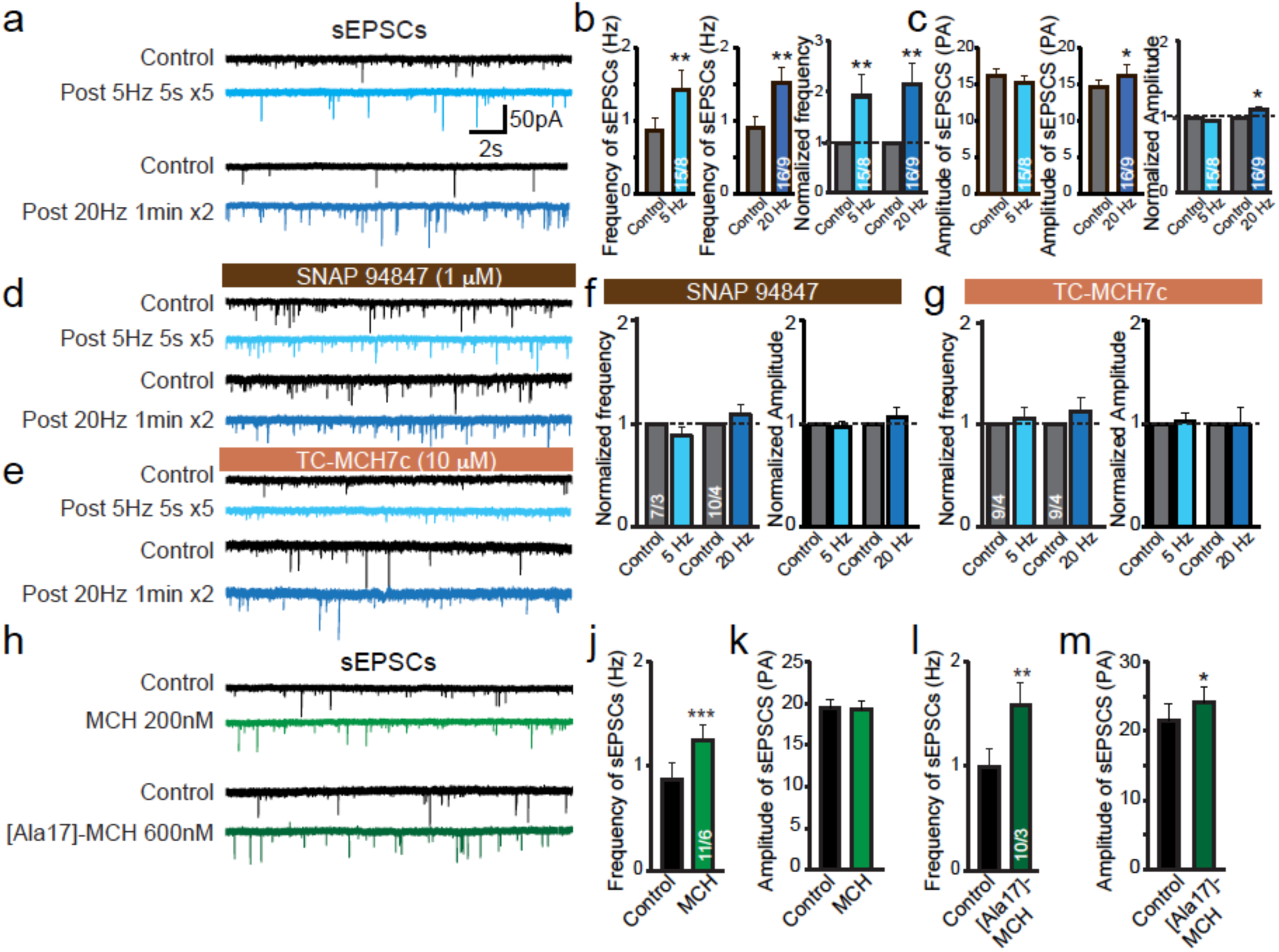
MCH enhances excitatory transmission via a presynaptic mechanism in the dLS-to-LHA projection neurons. **a**, Sample traces of sEPSCs before and after trains of optogenetic activation of MCH fibers. **b**, Pooled data of the comparisons of frequencies of sEPSCs. **c**, Pooled data of the comparisons of amplitudes of sEPSCs. **d&e**, Sample traces of sEPSCs before and after optogenetic activation of MCH fibers in the presence of SNAP94847 (d) and Tc-MCH7c (e). **f&g**, Normalized pooled data. **h**, Impact of exogenously applied MCH and [Ala17]-MCH on sEPSCs. **j-m**, Pooled data. Data are mean ± SEM; numbers of neurons/animals analyzed are indicated. Paired two-tailed Student’s t-tests were used: * p<0.05, ** p<0.01, ***p<0.001.

**Supplemental Figure 3.**
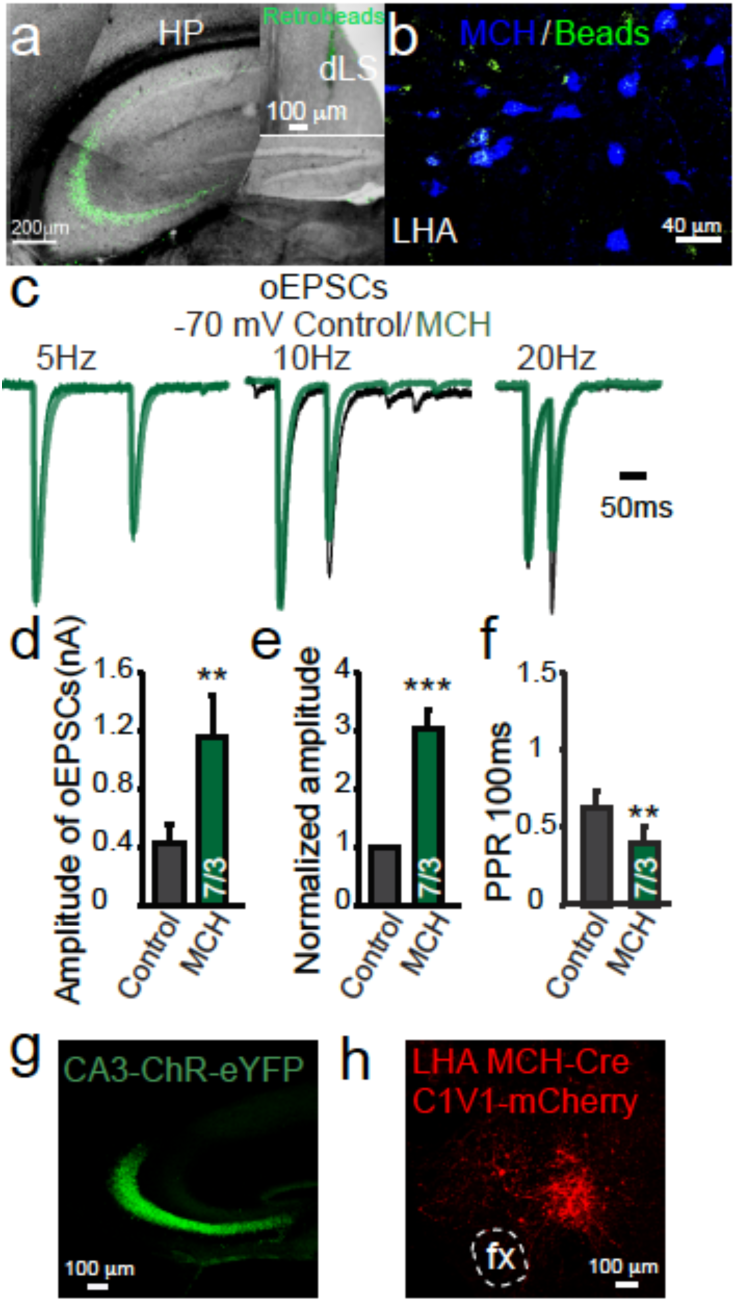
Exogenous MCH facilitates hippocampal excitatory inputs to dLS neurons. **a**, Sample images of retrograde labeled CA2/3 neurons after microfluorescent RetroBeads were injected into the dLS region. **b**, Retrograde labelled LHA neurons with immunoreactivity of MCH. **c**, Sample oEPSCs evoked by paired-pulse protocols at different intervals before and after the application of MCH. **d-f**, Quantification of oEPSCs originating from hippocampus in dLS neurons before and after the application of MCH. **g-h**, Sample images of injection sites. Data are mean ± SEM; numbers of neurons/animals analyzed are indicated. Paired two-tailed Student’s t-tests were used: * p<0.05, ** p<0.01, ***p<0.001.

**Supplemental Figure 4.**
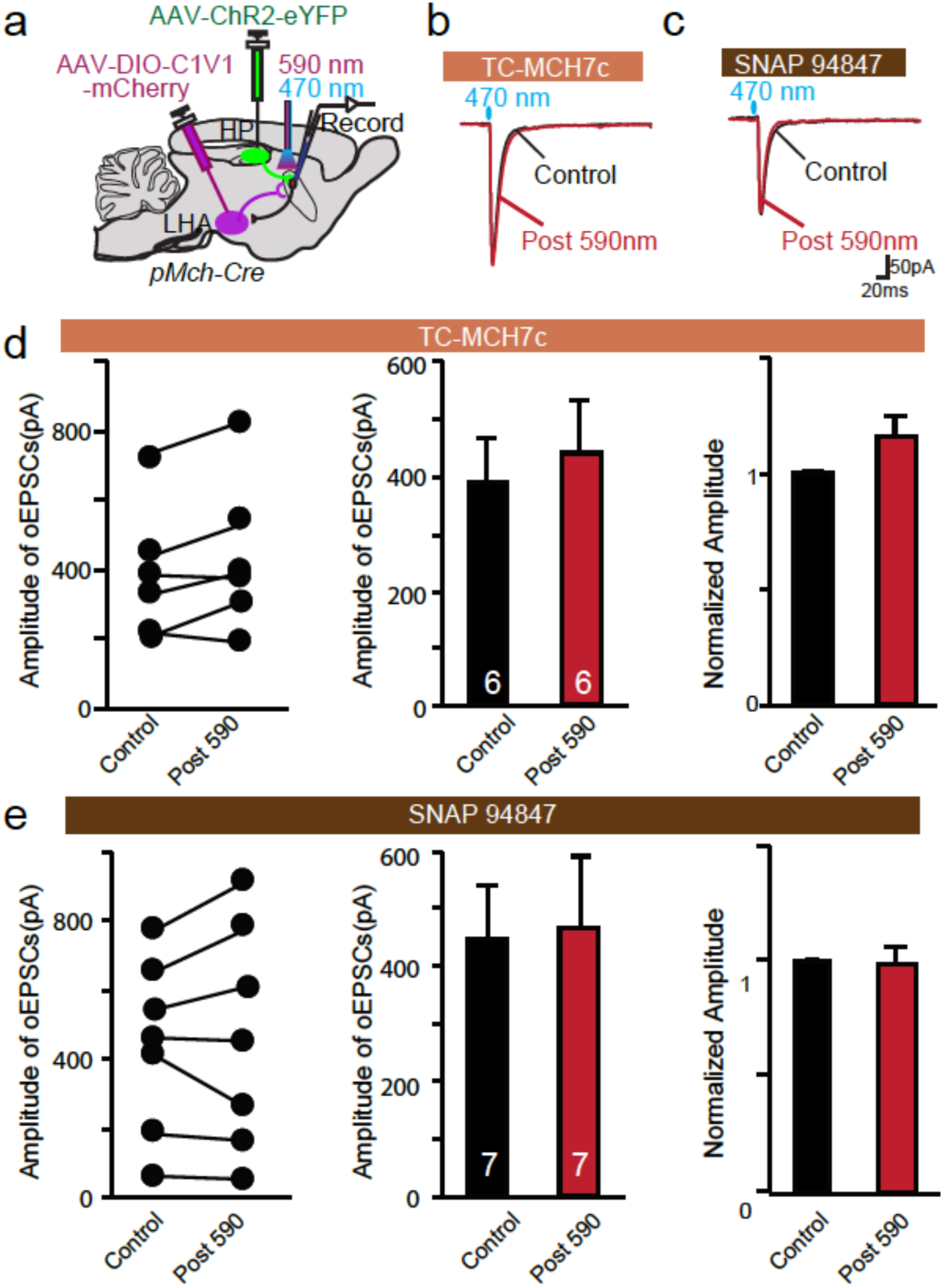
Endogenous MCH facilitates HP-to-dLS EPSCs via MCHR1 signaling. **a**, Diagram representing the experimental setup of dual optogenetic stimulations of hippocampal origin for oEPSCs by ChR2 (470nm) and of MCH neuronal origin for endogenous release of MCH peptide by red shifted C1V1ChR2 (590nm). AAV-ChR2-eYFP was injected into the ventral HP, while AAV-DIO-C1V1-mCherry was injected into the LHA of *p*MCH-Cre mice. dLS neurons were analyzed by whole cell recording. **b&c**, Representative superimposed traces of optogenetics-induced EPSCs in the presence of Tc-MCH7c (10 µM) and SNAP94847 (20 µM), in the dLS pre and post prolonged 590nm light activation on C1V1 positive MCH axons. **d&e**, Pooled data of amplitudes of single oEPSCs. Data are mean ± SEM; numbers of neurons analyzed are indicated in bars.

**Supplemental Figure 5.**
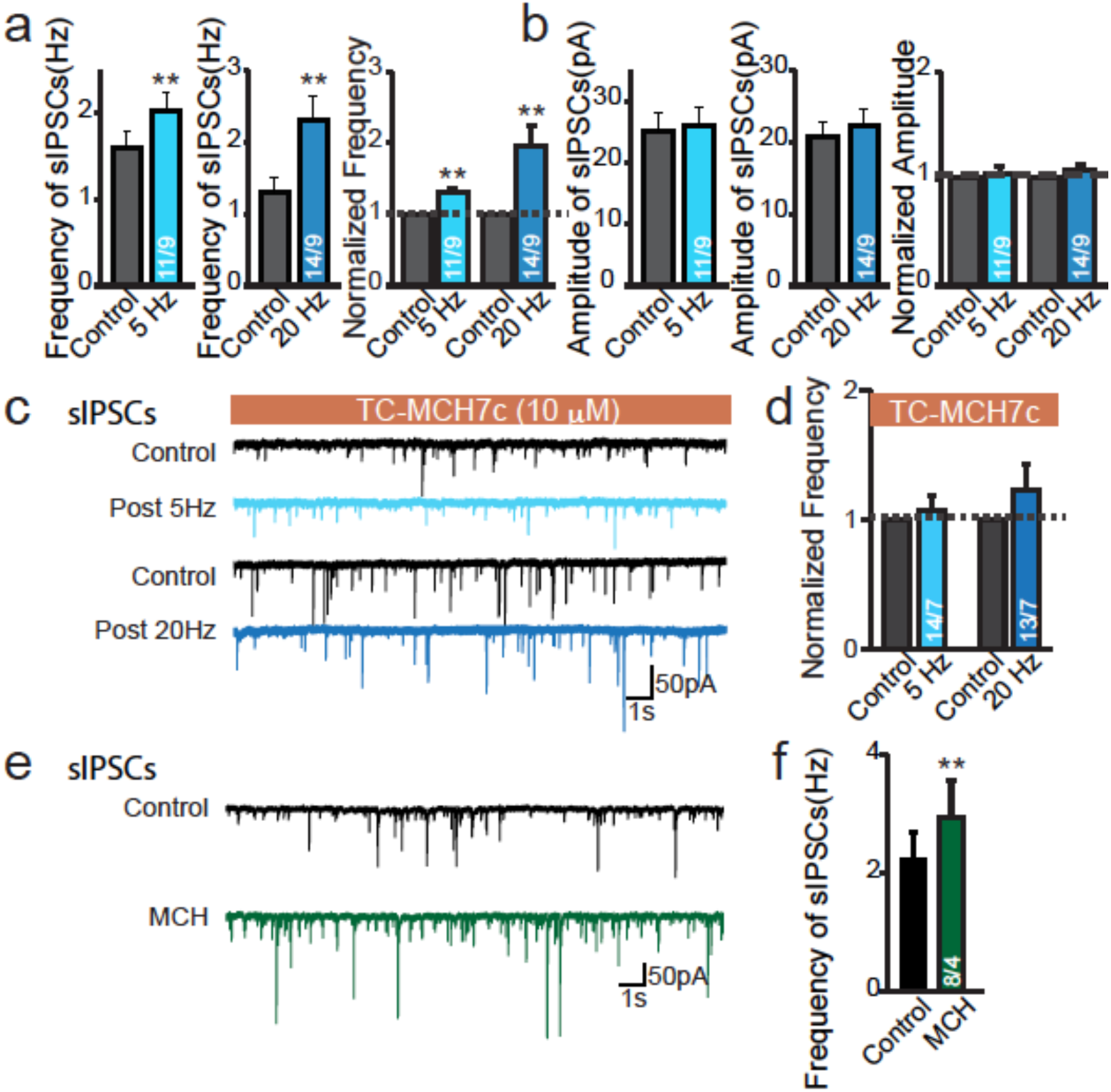
Endogenous MCH increases GABA tone in the dLS. **a**, Pooled data showing the changes of frequency of sIPSCs in dLS-LHA neurons, pre and post prolonged optogenetic stimulation of ChR-positive MCH axons. **b**, Pooled data for amplitude. **c**, Sample traces showing the impact of optogenetic stimulation of MCH fibers on sIPSCs in dLS neurons in the presence of Tc-MCH7c. **d**, Pooled data. **e**, Sample traces of sIPSCs before or after the application of MCH. **f**, Pooled data. Data are mean ± SEM; numbers of neurons/animals analyzed are indicated. Paired two-tailed Student’s t-tests were used: * p<0.05, ** p<0.01,***p<0.001.

**Supplemental Figure 6.**
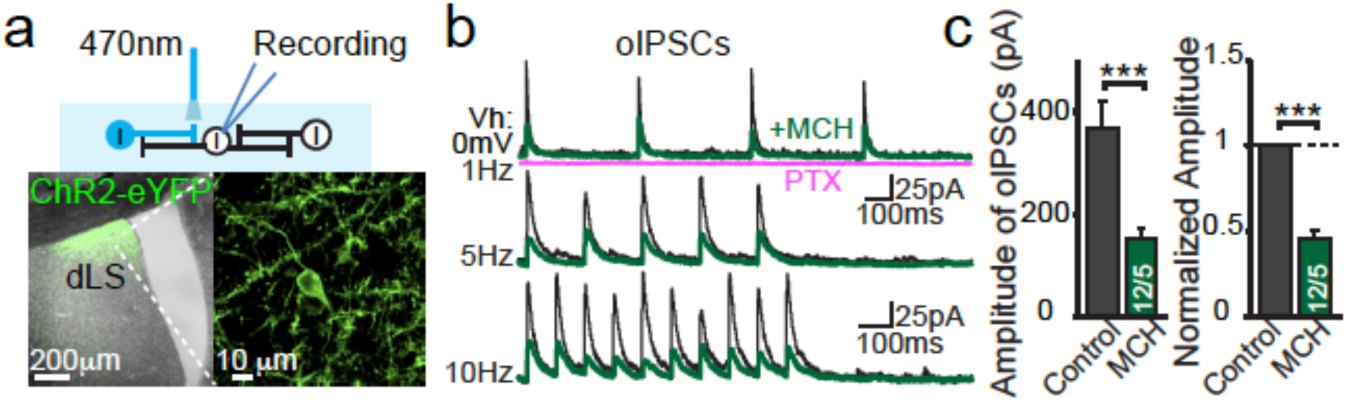
MCH suppresses collateral transmission in the dLS. **a**, Diagram of experimental setup and images showing ChR2 infection in the dLS. **b**, Sample traces showing the GABA_A_R-mediated inhibitory collateral synaptic transmission before (black traces) and after (green traces) administration of exogenous MCH neuropeptide (600nM). **c**, Pooled data. Data are mean ± SEM; numbers of neurons/animals analyzed are indicated. Paired two-tailed Student’s t-tests were used: ***p<0.001.

**Supplemental Figure 7.**
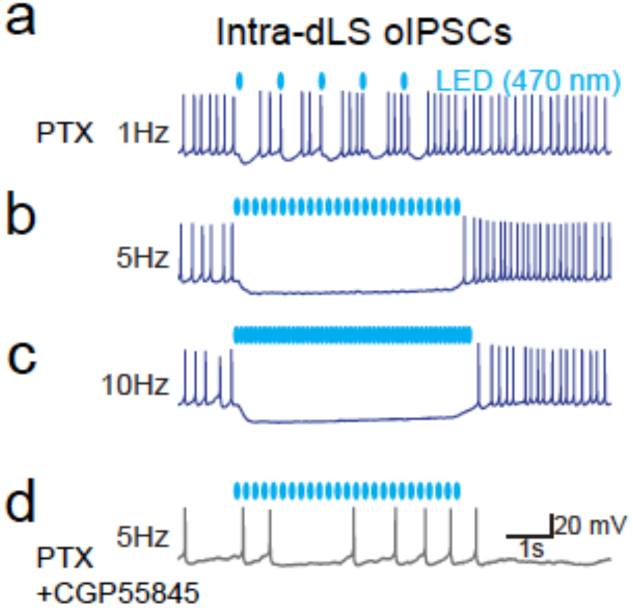
GABA_B_ receptor mediated inhibition suppresses spontaneous firing of dLS neurons. Sample traces showing the GABA_B_ receptor-mediated inhibitory synaptic transmission is sufficient to suppress dLS spontaneous action potential firing.

**Supplemental Figure 8.**
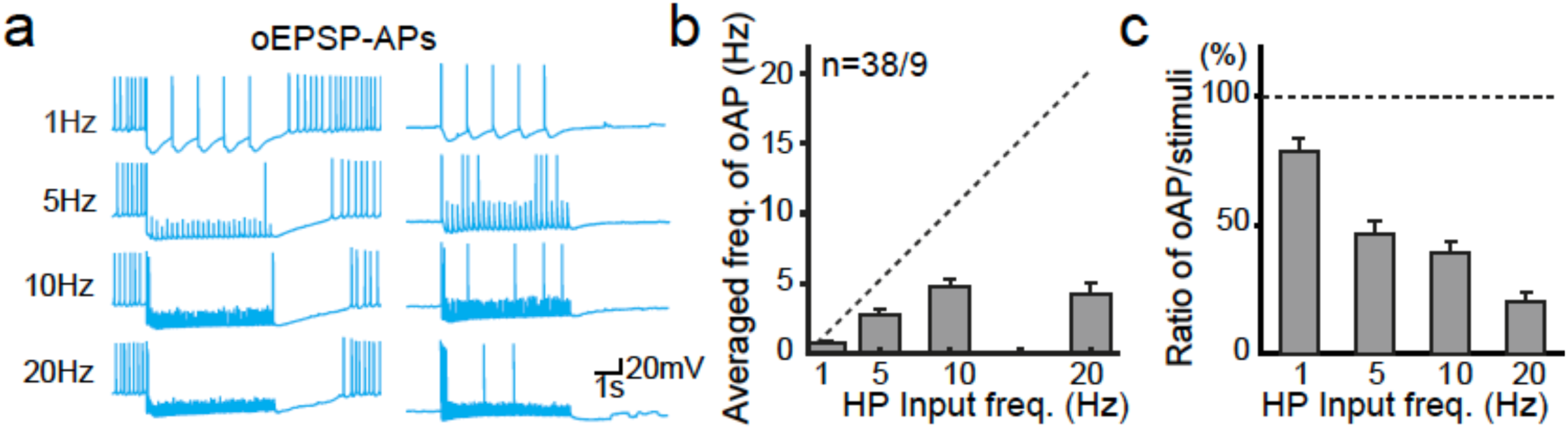
GABA_B_R-mediated FFI functions as a low pass filter. The FFI acts as a gate to block the high-frequency-encoded information generated in the hippocampus from passing through to downstream targets through the dLS neural network. Data are presented as mean ± SEM.

**Supplemental Figure 9.**
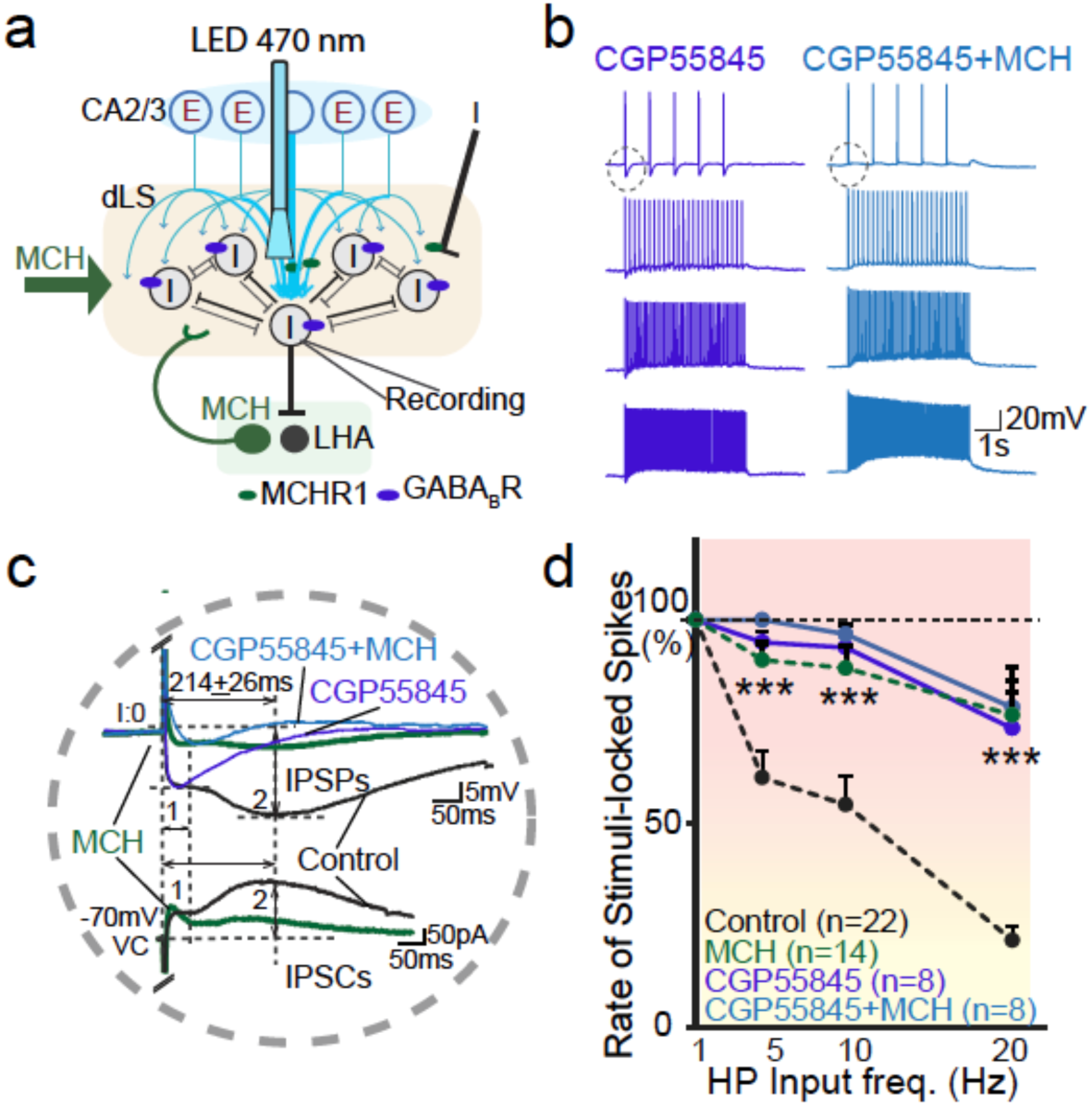
Hippocampal inputs induced postsynaptic action potentials. **a**, Possible wiring diagram of HP-dLS-LHA neurocircuitry. **b**, Sample traces of oEPSP-evoked action potentials in the presence of GABA_B_ blocker CGP55845 and MCH. **c**, Superimposed postsynaptic potentials (*upper panel*) and postsynaptic currents (*lower panel*) under different conditions. **d**, Pooled data depicts of the number of oEPSP-APs of dLS neurons as functions of the number of hippocampal optogenetic stimulations. Control and MCH data shown in Fig. 4 are shown here using dashed lines for comparisons. Data are presented as mean ± SEM; Two-tailed Student’s t-tests as well as ANOVA tests were used: * p<0.05, ** p<0.01, ***p<0.001.

**Supplemental Figure 10.**
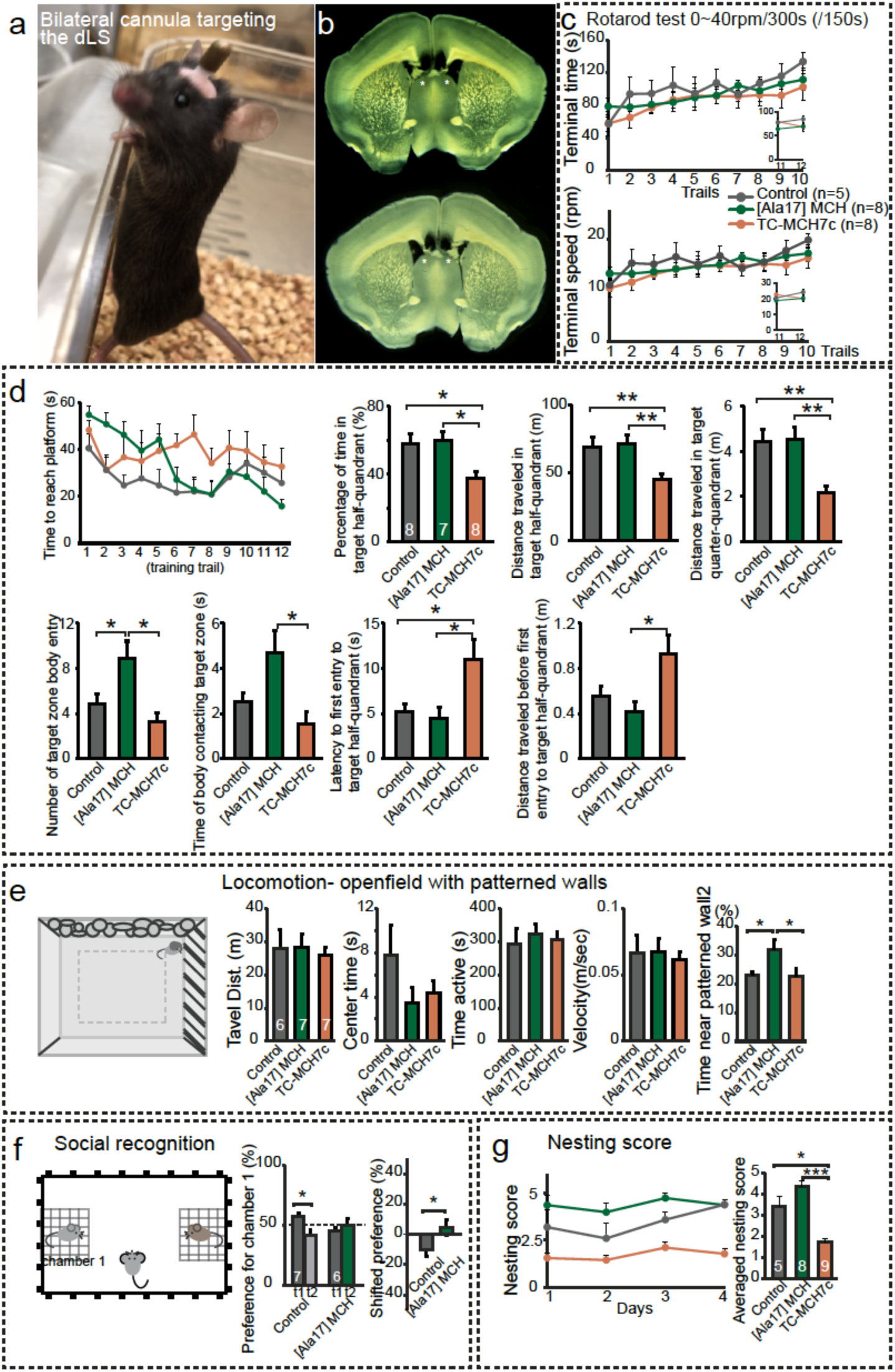
Behavioral impacts of MCH signaling in the dLS. **a**, Bilateral cannula implanted animal. **b**, Sample images showing the cannulae sites. **c**, Motor leaning in RotaRod system. **d**, Parameters for Morris water maze analyses in animals. **e**, Locomotor behavioral analyses. **f**, Social recognition. **g**, Nesting behavioral analyses. Data are presented as mean ± SEM; numbers of animals analyzed are indicated in bars. Two-tailed Student’s t-tests as well as ANOVA tests were used: * p<0.05, ** p<0.01, ***p<0.001.

